# *Anopheles stephensi* larval habitat superproductivity and its relevance for larval source management in Africa

**DOI:** 10.1101/2025.01.23.633752

**Authors:** Solomon Yared, Dereje Dengela, Peter Mumba, Sheleme Chibsa, Sarah Zohdy, Seth R Irish, Melissa Yoshimizu, Meshesha Balkew, Albert Akuno, T. Alex Perkins, Gonzalo M Vazquez-Prokopec

**Author notes:** **Gonzalo M. Vazquez-Prokopec Email:**. **Author Contributions:** <u>Designed research</u>: SZ, SI, MY, MB, SY. <u>Performed research</u>: SY, DD, PM, SC, MB, AA. <u>Analyzed data</u>: AA, TAP, GVP. <u>Wrote the paper</u>: SY, GVP, AA, TAP. **Data Availability:** All data and code will be made available upon publication from the following site: Vazquez Prokopec, Gonzalo, 2024, “Anopheles stephensi larval habitat superproductivity and its relevance for larval source management in Africa”, https://doi.org/10.15139/S3/NMP3LC, UNC Dataverse, DRAFT VERSION. **Competing Interest Statement:** Nothing to declare.

## Abstract

The invasion of Africa by *Anopheles stephensi* poses a significant threat to malaria elimination. As *An. stephensi* exploits a wide array of urban artificial larval habitats, it may be less impacted by rainfall variability compared to other native *Anopheles* species. We empirically investigated this assumption by quantifying the seasonal transition of an established population from eastern Ethiopia between rainy and dry periods. Monthly larval surveys generated evidence of significant heterogeneity between seasons in the type of habitat and their productivity. As the dry season progressed, *An. stephensi* productivity significantly concentrated in large water reservoirs (for drinking and construction) to a point in which up to 77% of all larvae originated from 23% of the sites. Such superproductive sites were primarily water cisterns used for residential or construction purposes. A two-patch metapopulation model of *An. stephensi* linked to rainfall data recreated the seasonal larval dynamics observed in the field and predicted that larval control targeted on superproducer water reservoirs, when implemented at coverages higher than 60%, may lead to *An. stephensi* elimination. Our findings highlight the role of environmental variability in regulating *An. stephensi* populations and open the window for the deployment of control strategies that exploit major mosquito population bottlenecks.

## Introduction

As the mosquito *Anopheles stephensi* continues its invasive range expansion within Africa (2), major questions about its ecology, bionomics, and epidemiological role still remain (3). In its native range (centered in India, Iran, Pakistan, and the Arabian Peninsula) *An. stephensi* is a major malaria vector found to be highly competent to both *Plasmodium falciparum* and *Plasmodium vivax (4)*. It is estimated that 12-15% of all urban malaria cases in India are vectored by *An. stephensi* (5). In Africa, the rapid increase in malaria case reports in Djibouti between 2017-2020, five years after the first detection of *An. stephensi*, (6) sent an alarming message about the potential for increased malaria incidence in the context of major and sustained reductions in morbidity and mortality throughout the continent (7-9). Observational studies suggest increased malaria rates are beginning to occur within Ethiopia, however, there seems to be heterogeneity in urban malaria increases in sites where *An. stephensi* has been detected *(4, 10-12)*. A challenge with assessing the epidemiological role of *An. stephensi* is often due to the limited knowledge of human-mosquito contacts, which depend on the level of productivity of larval habitats near human habitations and the flight and resting behavior of mosquitoes.

In its native habitat, *An. stephensi* exists as three biological forms that seem to be separated by environmental factors (the ‘type’ form is highly adapted to urban centers and feeds on humans, the intermediate form is found in peri-urban settings, and the ‘mysorensis’ form inhabits rural habitats and tends to feed more frequently on animals(13)). Reports from Africa indicate that the species is primarily urban, although non-urban detections have also been seen (14). In fact, urbanization itself appears to be favoring *An. stephensi* establishment primarily via the availability of large and permanent habitats in construction sites due to water storage associated with brick curing (15). Interestingly, reports of larval habitats for *An. stephensi* in Ethiopia have either focused on cross-sectional characterizations during the dry season (15) or rainy season (16-20). Unlike native *Anopheles* vectors, the ability for *An. stephensi* to persist in artificial habitats creates opportunities for populations to survive and thrive throughout the prolonged dry season of Ethiopia. Therefore, understanding the seasonal transition between rainy and dry periods is critical to identify better approaches for *An. stephensi* control.

Larval source management (LSM) combines a suite of methods that aim to prevent the development of mosquito immature stages, such as habitat modification/manipulation (reducing the availability of larval habitats) or larviciding/biological control (adding substances or organisms that either kill or inhibit the development of larvae) (21). Particularly for malaria prevention, LSM has shown highly variable levels of impact (22), and WHO recommends that larviciding only be implemented in areas where larval habitats are few, fixed and findable (21). In India, larval control of key *An. stephensi* larval habitats such as rice paddies and artificial containers has led to important reductions in *Plasmodium* infection in mosquitoes and humans (23). However, the challenges of implementing LSM in African cities invaded by *An. stephensi* are multiple and include largely unknown distribution of larval habitats within cities, the extent of coverage required for covering large urban centers, and limited evidence that methods that worked in the Asian context would work in Africa.

The persistence of *An. stephensi* in its invasive range is likely influenced by a complex interplay of factors, including the availability of suitable larval habitats, favorable climatic conditions, and the abundance of blood sources. This study focused on the seasonal fluctuation in *An. stephensi* populations by investigating the transition in larval productivity between the rainy and dry seasons in an established Ethiopian population. To understand the implications of these seasonal population changes for vector control, we linked our observations to a mathematical model that accounts for variations in larval habitat productivity and rainfall. This model explored the potential of LSM during the dry season as a strategy for reducing or locally eliminating *An. stephensi* populations.

## Results

### Characterizing *An. stephensi* habitat productivity

A total of 90 water holding containers and potential larval habitats were identified from 100 residences surveyed from the town of Kebri Dehar (Somali Region, Ethiopia) in November 2020 (Table 1). The majority (75%) of habitats were ground level cisterns (a mix of large water reservoirs used for household needs or construction purposes). In those habitats, 13,635 *An. stephensi*, 1,168 *Ae. aegypti*, and 757 *Culex* spp. larvae were collected throughout the four survey months. Habitat positivity (the proportion of habitats with larvae of *An. stephensi* and/or *Ae. aegypti* present) varied throughout the transition of rainy to dry season (Figure 1A). For *An. stephensi*, positivity decreased from 61% (55/90) in November to 53% (38/72) in December, followed by an increase to 74% (52/70) in January and 82% (63/77) in February (Figure 1B).

**Table 1.** Description of each potential larval habitat type identified in 100 visited houses and followed monthly between November 2020 and February 2021.

| Row Labels | Count of Type | Mean distance to house (m) | Mean area (m<sup>2</sup>) | Mean volume (L) | Number with water (sampled) |  |  |  |
| --- | --- | --- | --- | --- | --- | --- | --- | --- |
|  |  |  |  |  | November | December | January | February |
| Ground level cisterns | 68 | 6 | 11 | 873 | 70 | 68 | 70 | 70 |
| Tires | 12 | 8 | 6 | 8 | 12 | 0 | 0 | 0 |
| Ground level barrels | 6 | 5 | 7 | 268 | 6 | 6 | 6 | 6 |
| Rain catchments | 1 | 15 | 4 | 20 | 1 | 0 | 0 | 0 |
| Car wash drainage | 1 | 3 | 2 | 40 | 1 | 1 | 1 | 1 |
| Total | 90 | 7 | 10 | 683 | 90 | 75 | 77 | 77 |

**Figure 1.**
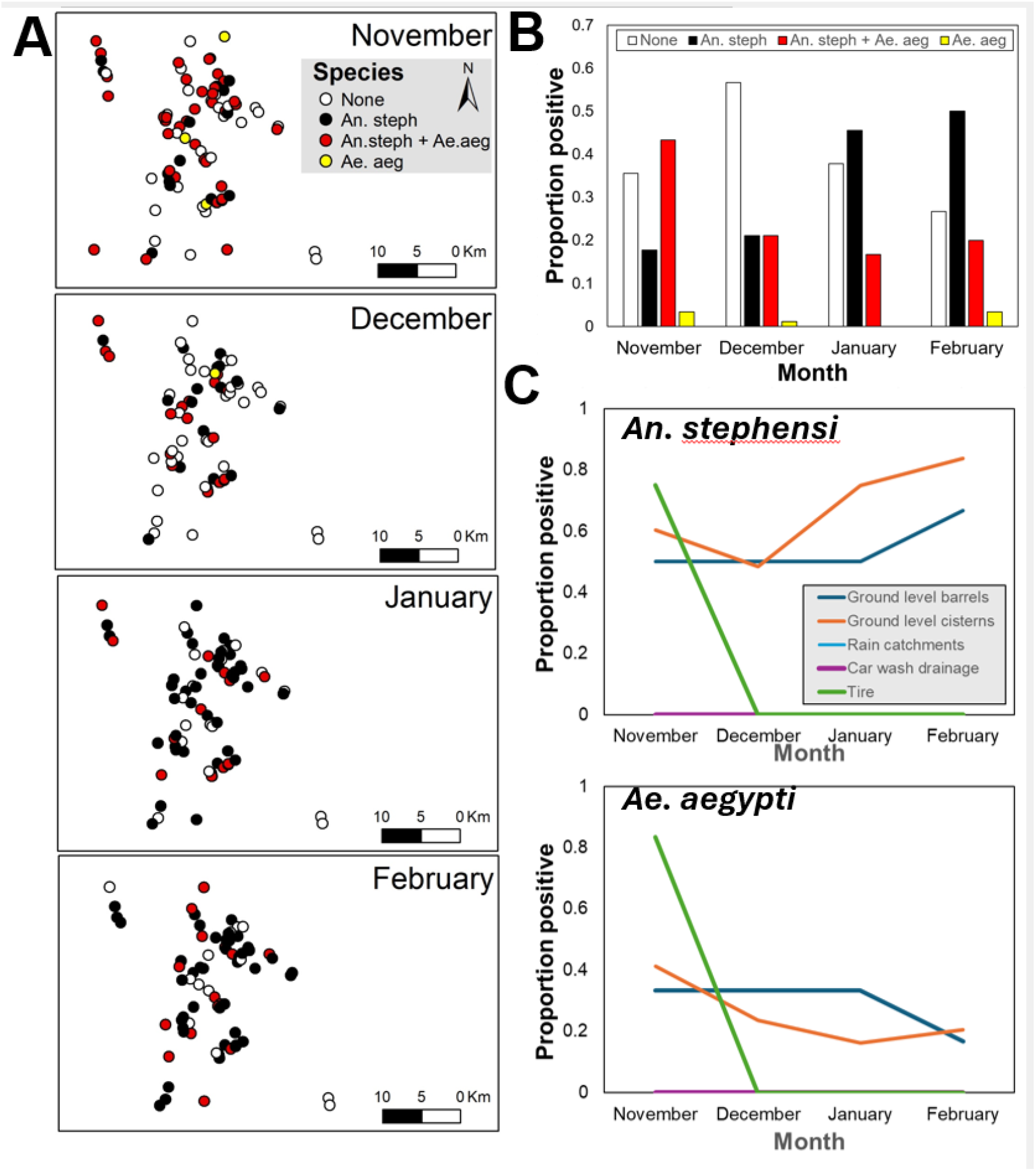
Seasonal transition in positivity of *An. stephensi* and *Ae. aegypti* in Kebri Dehar, Ethiopia. (A) Map of sites positive by each species alone or together throughout the seasonal shift. (B) Positivity (measured as proportion of habitats with either species alone or together) throughout the seasonal shift. (C) Positivity by type of larval habitat for each species.

Conversely, *Ae. aegypti* positivity reduced and maintained reduction from 47% in November to 28% in December, 20% in January and 21% in February (Figure 1B). Changes in positivity by both species were habitat- and species-specific (Figure 1C). Whereas for *An. stephensi* the seasonal increase in positivity was primarily due to increase in the number of positive cisterns and construction pits (Figure 1C), for *Ae. aegypti* the marked seasonal reduction was driven by a large number of tires and small larval habitats drying-up (Figure 1C).

*Anopheles stephensi* larval productivity (number of larvae per 20-dips per site) was highly heterogeneous (Figure 2A) and was significantly predicted by a negative binomial distribution on all months (Figure S6). When exploring the parameter K of the negative binomial (which measures the extent of heterogeneity in the data), evidence of extreme overdispersion (parameter *k*<0.5) in *An. stephensi* productivity was observed on all months (Figure S6A).

**Figure 2.**
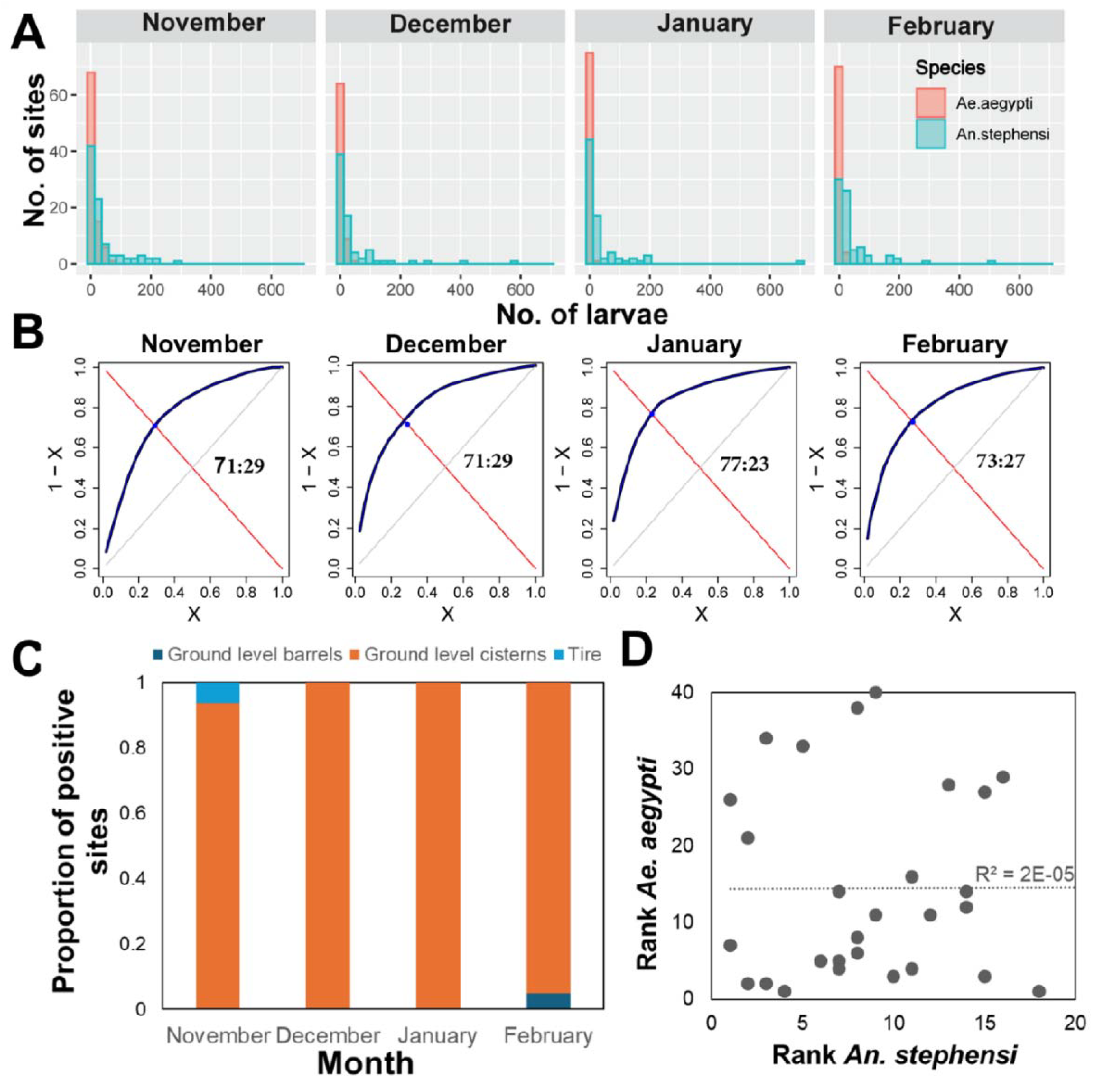
Heterogeneity in *An. stephensi* and *Ae. aegypti* larval productivity. (A) Histogram of the frequency of sites with a given number of larvae per 20 dips. (B) Results of the Pareto function applied to the number of *An. stephensi* per positive habitat. Numbers inside each panel indicate the pareto fraction (% of larvae : collected in % of larval sites). (C) For the top ith% larval sites, the type of habitat where An. stephensi was collected. (D) Rank of the top *An. stephensi* most productive larval habitats as a function of the top *Ae. aegypti* most productive habitats. The line indicates the result of a linear regression fit, with R^2^ shown inside the figure.

The dramatic heterogeneity in larval productivity observed for *An. stephensi* was further studied by calculating the Pareto fraction (Figure 2B). Overall, significant heterogeneity was detected across all months. In November, 29% of containers contributed 71% of *An. stephensi* larvae (71:29 in Figure 2B). Heterogeneity increased slightly as the dry season progressed, from 71:29 in November and December to 77:23 in January and 73:27 in February (Figure 2B). All such values indicate that most of the *An. stephensi* larvae concentrate in a few larval sites, with ground level cisterns as the primary habitat where *An. stephensi* concentrates (Figure 2C). Comparing the ranking of superproductive sites (in decreasing order of larval productivity) between species led to the conclusion that *An. stephensi* and *Ae. aegypti* primary producing sites do not overlap (Figure 2D). A large proportion (7/11, 64%) of *An. stephensi* superproductive sites in November were found to continue being superproducers on at least one of the subsequent months (Figure S7). No larval site was found to be superproductive throughout all months, and only 5 (13%) sites were found to be superproductive in three out of the four months.

### Implications of heterogeneous productivity for LSM

The Kebri Dehar findings imply the existence of two types of habitat patches for *An. stephensi* larval persistence: stable habitats that remain with water and produce large numbers of larvae throughout the year (e.g., cisterns and construction pits) and ephemeral habitats that either dry-up or lower their productivity as the dry season progresses (e.g., tires, small buckets, ground-level barrels). Modeling *An. stephensi* population dynamics using default parameters (Table S1 and Kebri Dehar rainfall) in both habitat patches reproduced the similar magnitude difference in larval productivity observed between them in the field as well as clear seasonality in adult abundance driven by rainfall (Figure 3A).

**Figure 3.**
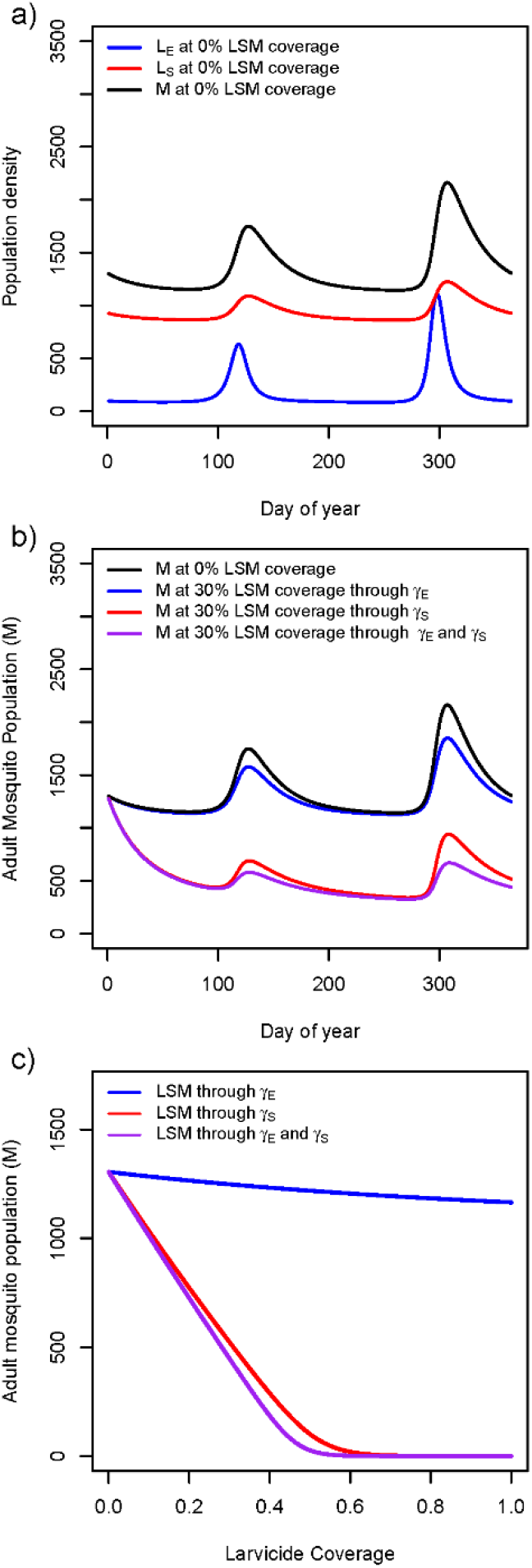
Numeric results of an *An. stephensi* population model that incorporates seasonality, two-patch dynamics, and the impact of larval control. (a) The dynamic of mosquito populations in the ephemeral (E) and stable (S) patches for both larvae (L_i_) and adult mosquitoes (M) when no control is implemented. (b) Simulating the impact of larval control at 30% coverage in the E, S and E+S patches on the number of adult mosquitoes in the entire population. (c) impact of larval control on the adult mosquito population as a function of intervention coverage for interventions targeted at the E, S and E+S patches.

Simulating the implementation of residual larviciding during the dry season (February) with a larvicide that lasts up to six months and at a coverage of 30% led to a minor reduction in *An. stephensi* adult density when the ephemeral patch was treated (increase in γ_E,_ Figure 3B). When the stable patch was treated at the same coverage and with the same larvicide, a dramatic reduction in adult mosquito population was observed (increase in γ_S_, Figure 3B). Interestingly, simultaneously applying LSM on both the stable and ephemeral patches (increase γ_S_ and γ_E_) led to a very similar reduction in adult mosquitoes than controlling the stable patch alone (Figure 3B). Each intervention strategy showed a different impact on the adult population as the intervention coverage increased (Figure 3C). Controlling the ephemeral patch led to minor impact on adult mosquitoes (∼10% reduction). When the intervention was conducted on the stable patch, adult mosquito population size was exponentially reduced, with vector elimination occurring at LSM coverages above 60% (Figure 3C). Interestingly, controlling both ephemeral and stable patches led to vector elimination at 50% coverage of all habitats, a minor vector control gain compared to targeting the stable patch alone (Figure 3C).

## Discussion

*Anopheles stephensi* invasion of Africa has led to significant challenges for local and national malaria control programs aiming at containing this urban container-breeding mosquito. Our study provides important information about *An. stephensi* larval ecology and the potential for vector elimination through data-informed and targeted LSM. In the highly seasonal urban centers of Ethiopia, *An. stephensi* experience a major population bottleneck during the dry season, concentrating productivity in large water cisterns associated with residential water consumption or construction. The stability of nutrient-rich water throughout the year makes cisterns and construction pits ideal productive habitats, leading some of them to become superproductive (accounting for up to 70% of all larvae collected). Targeting LSM with a long-lasting larvicide on such stable and superproductive habitats at a coverage above 60% is predicted to lead to major population reductions and vector elimination even if ephemerous habitats are left untreated. Our findings can have profound implications for the design of LSM plans against *An. stephensi* in Africa and beyond.

During the rainy season, *An. stephensi* is commonly found in a wide array of larval sites throughout Ethiopia (18). Conversely, during the peak of the dry season in Jigjiga (Somali Region) *An. stephensi* was found infesting a few locations, primarily ground level cisterns associated with construction (15). Unifying such findings, here we show that *An. stephensi* undergoes a major population transition between the rainy and dry seasons, concentrating in ground level cisterns and disappearing from smaller artificial containers due to them drying out. While positivity was informative for identifying the array of potential larval habitats, larval productivity became more relevant for identifying the potential contribution of each site to the maintenance of *An. stephensi* populations. We found significant levels of heterogeneity in *An. stephensi* larval productivity, evidenced by the contribution of up to 77% of all larvae by only 23% of the larval sites. The trend of aggregation was consistent across months, with an observed increase in the level of heterogeneity as Kebri Dehar transitioned from the rainy to the dry season. Unfortunately, *An. stephensi* adult collection numbers in Ethiopia tend to be very low and centered in peridomestic habitats (18, 24), limiting our ability to correlate superproductivity with adult mosquito density. While a correlation between heterogeneity in *An. stephensi* adult density and malaria transmission potential are likely (as observed for *An. gambiae* in Uganda (25)), future studies should focus on superproductive habitats and their potential role as sources for adult *An. stephensi* and increased risk of urban malaria transmission.

In Africa, *Ae. aegypti* has been reported anecdotally cohabiting with *An. stephensi* in various larval sites (14, 18, 26). Here we show that the level of overlap varies with the season and the type of habitat. During the rainy season, both species exploit similar containers (plastic containers and tires) and can be found in other habitats not surveyed in Kebri Dehar (19). As the dry season progresses, the lack of rain coupled with high heat leads to rapid water evaporation in small containers and tires and to the disappearance of both mosquito species from them. It is possible that *Ae. aegypti* survives the long dry season due to the large crop of eggs remaining in dried-out plastic containers and tires. As water holding containers remaining in the dry season are primarily ground cisterns and large plastic drums, it is possible that *Ae. aegypti* faces additional sources of mortality due to other mosquito larvae, predators and increased sun irradiation in open cisterns and construction pits. Combined, such factors may be responsible for the low productivity of *Ae. aegypti* and the high productivity of habitats by *An. stephensi*. More research on the larval ecology of both species is needed to understand whether cohabiting can be used to leverage LSM implementation. Our findings suggest that the most productive habitats for *An. stephensi* are not necessarily the most productive habitats for *Ae. aegypti*, challenging the idea of efficient control of both species if specific habitats are targeted.

We acknowledge several limitations of our study. First, we focused our sampling on 100 houses with 90 habitats and it is likely we missed specific larval sites that have been described by others as infested by *An. stephensi* during the rainy season (e.g., buckets and small plastic containers). Given what we have seen with tires and other containers, those small habitats are very likely prone to drying and may not contribute to significant productivity in the dry season. Second, pupae are often used to quantify productivity. Unfortunately, our study only detected pupae in a few instances (80 sites out of 319 with water) and the field team was unable to identify them to species. Therefore, more complex measures of productivity (pupae per dip, pupae per person (27)) were not attempted and should be the focus of future research.

Considering that *An. stephensi* eggs only survive desiccation for a few days or up to two weeks (28) the finding of increased heterogeneity in the dry season could be of significant relevance for LSM deployment. Our empirical study and modeling findings imply that *An. stephensi* populations may be shaped by seasonally-driven metapopulation dynamics. While in the rainy season population mixing is high and occurs across both stable (e.g., large cisterns, construction pits and other large water reservoirs) and ephemeral (e.g., tires, small containers and other water reservoirs), in the dry season stable habitats which are replenished by filling tap water from time to time function as sources of adult mosquitoes that maintain the population throughout the dry season. Given this strong seasonality in productivity, our model suggest that targeting larval control in stable habitats during the dry season can lead to significant population reductions and even vector elimination if coverage is above 60%. Our model also shows how sensitive findings may be to the efficacy of the larvicide. As larvicides such as pyriproxyfen (29) and spinosad (30), which have long-lasting formulations that last up to six months, continue being evaluated and showing high efficacy against *An. stephensi*, the opportunity for sustained long-term control becomes more attainable. Other forms of control, including mosquito proofing water cisterns (31) or the introduction of larvivorous fish (32) may be biorational alternatives to the use of larvicides, particularly in stable habitats that remain with water for years. Our findings, if validated empirically with field trials, suggest that the goal of *An. stephensi* elimination from Ethiopian cities with precision LSM at adequate coverage may be within reach.

## Materials and Methods

### Study design

We studied a well-established population of *An. stephensi* in the semi-arid town of Kebri Dehar, Somali Region, Ethiopia, where the species was first detected in 2016 (SI Text and Figure S1). Since its introduction, *An. stephensi* has expanded throughout the city and has been indirectly implicated in increased malaria transmission (12). Kebri Dehar is located in a semi-arid area that receives only an average of 200 mm of rain per year, distributed in two rainy periods (Figure S3). The temperature and rainfall for the period of study are shown in Figure S2.

To characterize water presence and mosquito productivity in a cohort of containers with potential to be productive during the rainy season and through the transition between rain (November, December) and dry (January-February) periods we initially surveyed 100 residences from Kebri Dehar in November 2020. All containers with water in November 2020 were geocoded (GPS Garmin eTrex) and monitored monthly until February 2021. During each survey, a habitat was first visually inspected for the presence of water. When water was present, 20 standard dips were taken (350ml capacity) from each identified habitat by sampling its entire area. Distance to the nearest house, vegetation coverage, volume of water in the containers and habitat type (i.e., ground level cisterns, ground level barrels, plastic containers, car wash drains and rain catchments) were recorded. The *Anopheles* larvae were separated from the culicine larvae based on morphology and allowed to emerge to adult in cages separated by habitat for species identification using a dissecting microscope and a standard keys (33).

### Statistical Analysis

Larval productivity was quantified as the total number of larvae present in the 20 dips per habitat and month. Heterogeneity in habitat productivity was quantified in two ways. A negative binomial fit to the distribution of larvae per container (per 20-dips) was used to estimate (via Maximum Likelihood) the parameter k, which is a measure of overdispersion in the data (34). Values of k<0.5 indicate strong overdispersion (34), meaning most of the larvae are aggregated (i.e. found in a few containers). The Pareto fraction (25) was used to further quantify heterogeneity. We define Pareto fraction for the number of larvae per site (*N*_*Li*_) from its empirical CDF (eCDF) as Cooper et al. (25), with *X* denoting the proportion of *N* observations and *Y* the empirical cumulative distribution: 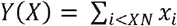 *Y(X)* monotonically increases with *X* and intersects the line 1-*X*. The point *X* where *Y(X)* = 1-*X* defines the Pareto fraction. At that point, there is a *X*:1-*X* rule, for instance if *X*=0.8 then the Pareto fraction corresponds to the 80-20 rule. The Pareto index for a given fraction can be calculated as 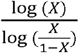 We calculated the Pareto fraction and Pareto index for each month. From the Pareto fraction, we identified the %^ith^ larval sites that contributed the most to total larval productivity (e.g., the top 20% of sites that contribute to 80% of larvae collected). Those ‘superproducer’ habitats were mapped to understand patterns of productivity across space and time. Additionally, ordinary least squares regression was used to correlate the rank of the most productive habitats (1 being most productive and least productive in ascending order) for *An. stephensi* and *Ae. aegypti*.

### Model of seasonal LSM in heterogeneous landscapes

The Kebri Dehar data was used to parameterize a multi-patch model that explicitly incorporates heterogeneity in larval productivity to explore how the seasonal transition in *An. stephensi* populations can be exploited to improve LSM. Here, we expanded the model developed by Smith et al. (35), which explicitly considers the effect of larval crowding and density-dependent competition within a multi-patch framework (see SI Text), by incorporating seasonality in rainfall and increased larval mortality due to loss of water in certain habitat types as the dry season progressed.

Specifically, we structured the model in patches, which can contain many aquatic habitats (called pools in (35)) of a given type such as construction pits, ground cement cisterns, water storage containers at home, etc. In this model *L*_*i*_(*t*) represents the number of larva in patch *i ∈*{1,2,…,*n*}, where *n*, is the total number of patches, and *M* represents the number of adult mosquitoes in the whole system of *n* patches, modeled together as one. The model, is formally structured as:

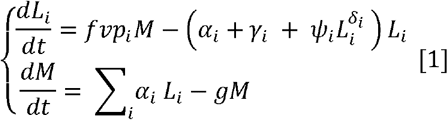

Where *i=*1,2,…,*n* is the index for any patch, the mosquito larva reside in *n* different patches, while the adult mosquito population is treated as a single unit. Other model parameters include mosquito blood feeding rate (*f*) the number of eggs laid by a mosquito each egg laying cycle (*v*) the per-capita death rate of adult mosquitoes (*g*) the fraction of eggs laid in Patch *i*(*p*_*i*_) the maturity rate of mosquitoes in Patch *i*(α_*i*_) the Patch *i* per-capita death rate not caused by overcrowding (γ_*i*_) the patch-specific increase in per-capita mortality in response to crowding (ψ_*i*_) and the Patch *i* mean crowding (δ_*i*_). The parameters contained in the model are described

For tractability and simplicity, we generated a two-patch model where Patch 1 is ephemeral (containing water during the rainy season but rapidly evaporating as the dry season progresses, denoted with subscript E) and Patch 2 is stable (containing water throughout rainy and dry season, denoted with subscript S). Seasonality was incorporated in the model by linking parameter γ_*i*_ to rainfall (described in SI Text and Figure S3). To incorporate the effects of LSM, we assumed that larvicides increased *γ*_*E*_, or *γ*_*S*_, depending on whether control was done targeting either or both patches. Intervention coverage (proportion of habitats within the patch that were treated) was also explicitly incorporated using the following equations:

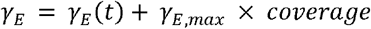

And

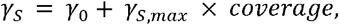

Where *γ*_*E,max*_*=kγ*_*E*_(*t*) and *γ*_*S,max*_*= kγ*_0_ The parameter *k* represents the **relative increase in larval mortality** that can be achieved when the larvicide is applied with full coverage (coverage = 1) compared to the baseline mortality rates *γ*_*E*_(*t*) and*γ*_0_which are the natural mortality rates of larva in the ephemeral and stable patches respectively, when no larvicide is applied (coverage =0). Then, *γ*_,*max*_ is the maximum additional patch-larval mortality that can be induced by larvicide application, and it is proportional to the natural or baseline patch-larval mortality rate (SI Text). For this study, we simulate LSM using *k*=20 and conduct sensitivity analyses of *k* to understand how larvicide efficacy impact results (Figure S5). To minimize the effects of the intervention on populations that were not yet in equilibrium, we ran the model for two years, implementing LSM in the beginning of the second year.

## Acknowledgments

Our gratitude extends to PMI VectorLink Ethiopia for their invaluable support in our field mosquito collection. Additionally, our appreciation goes to Yared and Ismail for their diligent efforts in collecting mosquito larvae from households. We are also thankful to the Kebridehar communities for their willingness to participate and for granting us access to inspect their water holding containers. Support for this study was provided by the US President’s Malaria Initiative. Gonzalo Vazquez-Prokopec was supported by the US Centers for Disease Control (CDC-BAA-75D301-23-R-72545). Albert Akuno and T. Alex Perkins were supported by a grant from the NIH National Institute of General Medical Sciences R35 MIRA program (R35GM143029).

## Disclaimer

The findings and conclusions expressed herein are those of the authors and do not necessarily represent the official position of the U.S. Centers for Disease Control and Prevention (CDC), U.S. Agency for International Development (USAID), or U.S. President’s Malaria Initiative (PMI).

## Notes

### Competing Interest Statement

The authors have declared no competing interest.

