## Supplementary material for "*Anopheles stephensi* larval habitat superproductivity and its relevance for larval source management in Africa": SI Text

Gonzalo M. Vazquez-Prokopec.

**This PDF file includes:**

Supporting text

Figures S1 to S7

Table S1

SI References

Supporting Information Text

**Study area and mosquito collections.** Kebri Dehar has a population of ~120,000 and is distanced 400 kilometers from Jigjiga City and 1,035 kilometers from the capital city of Addis Ababa. The majority of the study area's geography is lowland plains, with an average elevation of 525 meters above sea level and a few steeper foothills. The latitude and longitude of the area are 6° 44’25’’N/ 44°16’38’’E, respectively. There are few shrubs and trees in the area, including several *Acacia* species and incense trees. Kebri Dehar is a semi-arid area and the average annual temperature in the area is between 23 and 30 °C. The rainfall pattern in the area is bimodal, with peaks in May and November, and averages 200 mm. Despite these patterns, the rainfall is variable, making the area prone to recurrent droughts that can last several months.

Typically, each house contains an outdoor constructed cement tank used for storing water, referred to as a *birket* (Figure S1). Other typical structures used to harbor water are also shown in Figure S1. The house type of the study area is mainly rectangular with a corrugated iron roof and cement walls and often inhabited by multiple families.

The initial sampling involved any potential artificial larval breeding habitat with water such as ground level cisterns, ground level barrels, plastic containers, car wash drains and rain catchments (Figure S1). To reach the target number of 100 houses, the field team moved to the center of Kebri Dehar and began contacting random houses asking for permission to conduct an initial survey. If a water-holding container was found, homeowners were asked to consent to repeated surveys of their house for the months of December 2020, January 2021, and February 2021. Of the 100 houses visited, 15 did not have any potential habitats with water (tires, drums and cisterns were completely dry), leaving 85 houses and 90 habitats for the study. A total of 13,635 *An. stephensi*, 1,168 *Ae. aegypti* and 757 *Culex* spp. were collected throughout the four survey months in those sites. Given our inability to confirm whether *Culex* spp. were a single or multiple species, we only report *An. stephensi* and *Ae. aegypti*. A total of 13,068 (96%) larvae emerged as adults and were used to confirm the identity of each species from each container. The remaining 4% were only identified to Genus. Figure S1 shows the distribution of the 90 sampling sites, distinguished by habitat type, in Kebri Dehar whereas Figure S2 shows the trend in rainfall and temperature throughout the study period.

**Model parameterization**

In our proposed model, $L_{i}(t)$ represents the number of larva in patch $i \in\{1, 2, \cdot\cdot\cdot, n\}$, where $n$ is the total number of patches, and $M$ represents the number of adult mosquitoes in the whole system of $n$ patches, modeled together as one. That is, in the model, which is formally given as

$$\left\{ \begin{aligned} \frac{{dL}_{i}}{dt}=fvp_{i}M-\left( \alpha_{i}+\gamma_{i} + \psi_{i}L_{i}^{\delta_{i}} \right)L_{i} \\ \frac{dM}{dt}= \sum_{i} \alpha_{i} L_{i}-gM \end{aligned} \right. [1]$$

where $i=1,2,\ldots,n$ is the index for any patch, the mosquito larva reside in $n$ different patches, while the adult mosquito population is treated as a single unit. The parameters contained in the model are described in Table S1.

**Table S1.** Model parameters used to simulate *An. stephensi* population dynamics under seasonal and heterogeneous environments.

| **Parameters** | **Description** | **Default values** |
| --- | --- | --- |
| $f$ | Mosquito blood feeding rate | 0.30 |
| $v$ | Number of eggs laid by a mosquito each egg laying cycle | 25 |
| $g$ | The per-capita death rate of adult mosquitoes | 0.083 |
| $p_{i}$ | Fraction of eggs laid in Patch $i$ | 0.5 |
| $\alpha_{i}$ | Maturity rate of mosquitoes in Patch $i$ | 0.1 |
| $\gamma_{i}$ | Patch $i$ per-capita death rate not caused by overcrowding | 0.5 |
| $\psi_{i}$ | Patch-specific increase in per-capita mortality in response to crowding | 0.01 |
| $\delta_{i}$ | Patch $i$ mean crowding | 0.9 |

We structured the model to let some of the parameters be driven by the rainfall as an approach to capture the seasonality in mosquito production, as observed in the Kebri Dehar empirical study. To capture general variability in rainfall seasonality for Kebri Dehar, we used the daily averages of 2010 - 2020 rainfall data from the city’s airport (Figure S3). We then smoothed the raw data using quasi-Poisson family of generalized additive models (GAM) with integrated smoothness estimation (R package mgcv (1), Figure S3).

**Simulation of *An. stephensi* population dynamics**

For the simulation study, we partition Kebri Dehar into two patches, where we assume that Patch 1 is ephemeral, and Patch 2 is stable due to the presence of aquatic habitats that do not dry up during the dry season. To achieve the ephemerality of Patch 1, we let the larva mortality rate, $\gamma_{1}$, which we now denote as $\gamma_{E}$ (where the subscript $E$ stands for ephemeral), be driven by the Kebri Dehar daily rainfall data (Figure S3). In particular, we assume that $\gamma_{E}(t)$ is largest when rainfall on the day $t$ is lowest, and vice versa, signifying high larval mortality in this patch during the dry season, and a low mortality rate during the rainy season. For the lowest mortality rate, we use the default value in Table 1, and set the largest value to be 100 times the lowest value during the dry season (to differentiate productivity in different habitat types, as observed in the empirical study). To achieve this, and to incorporate the rainfall dynamics in the larval mortality rate, we use the relation:

$$\gamma_{E}\left( t \right)= \gamma_{0}+\left( 100\gamma_{0}- \gamma_{0} \right)\frac{\max\left( Y \right)-Y\left( t \right)}{\max\left( Y \right)-\min\left( Y \right)}[2]$$

where $\gamma_{0}$ is the base value, to which we assign the default γ value in Table 1, $Y$ is the rainfall data, and $t$ is time (day). In this equation, the value (100) that we use to multiply the lowest mortality rate is, in the current study, an empirical unknown, and we only defined it as a constant for illustrative purposes, as also described in the previous version of the model (2). This value, which we can refer to as the fold increase in the larval mortality rate in the ephemeral patch due to low precipitation, was invaluable in achieving the empirical 10-fold difference in larval density between the stable and ephemeral larval population during the dry season (see Figures S4a, S4b and S4c). In addition, there is no particular justification of relation (2), other than the fact that it allows for the incorporation of the rainfall data, and captures the desired dynamics of $\gamma_{E}$ being smallest on the day rainfall is highest, and vice versa. We thus have that the values of $\gamma_{E}(t)$ in the interval $[\gamma_{0}, 100\gamma_{0}]$ increases (decreases) with decrease (increase) in the daily rainfall data.

To make Patch 2 stable, we use constant parameters, and set its larval mortality equal to the ephemeral Patch 1 base larval mortality (i.e, we use $\gamma_{S}= \gamma_{0}$, where the subscript $S$ stands for stable), with the rest of Patch 2 parameters being constant as well. In addition, the rest of the ephemeral patch parameters are constant as well. This set-up captures the findings in the empirical study that there was stable mosquito productivity during the dry season due to the existence of large water reservoirs in Kebri Dehar. In this simulation study, we use the default model parameters from Smith et al. (2), with the exception of the patch-larval mortality rates (see Table 1).

**Modeling larval source management (LSM)**

We explored the options of controlling *An. stephensi* by simulating the implementation of LSM by assuming that a long-lasting larvicide was applied either in the ephemeral patch through $\gamma_{E}$, or in the stable patch through $\gamma_{S}$, or in both of the patches together through $\gamma_{E}$ and $\gamma_{S}$. During these simulations, the rain-driven mortality rate defined in Equation [2] is utilized as the base mortality rate in the ephemeral patch, while the base mortality rate in the stable patch is taken to be the default larval mortality value in Table 1. Then, the aim is to increase the patch mortality rates over and above these base values based on an array of increasing coverage values defined in the interval [0, 1]. Consequently, the mortality rates in the ephemeral and stable patches under larvicide coverage are respectively given as

$$\gamma_{E}= \gamma_{E}\left( t \right)+ \gamma_{E,max} \times coverage$$

and

$$\gamma_{S}= \gamma_{0}+ \gamma_{S,max} \times coverage,$$

where $\gamma_{E,max}=k \gamma_{E}\left( t \right)$ and $\gamma_{S,max}=k \gamma_{0}$. In this setting, the parameter $k$ represents the **relative increase in larval mortality** that can be achieved when the larvicide is applied with full coverage (coverage = 1) compared to the baseline mortality rates $\gamma_{E}\left( t \right)$ and $\gamma_{0}$, which are the natural mortality rates of larva in the ephemeral and stable patches respectively, when no larvicide is applied (coverage = 0), as defined by (2). Then, $\gamma_{. , max}$ is the maximum additional patch-larval mortality that can be induced by larvicide application, and it is proportional to the natural or baseline patch-larval mortality rate. This then implies that when $coverage=0$, no larvicide is applied, and for the ephemeral patch, $\gamma_{E}= \gamma_{E}\left( t \right)$, while for the stable patch, $\gamma_{S}= \gamma_{0}$, and when $coverage=1$ (full coverage), $\gamma_{E}=(1+k) \gamma_{E}\left( t \right)$ and $\gamma_{S}=(1+k) \gamma_{0}$, implying that at full coverage, the larval mortality rate is increased by a factor of $1+k$ compared to the baseline rate. Thus, a higher $k$ value ($k>1$) means that the larvicide is highly effective and can significantly increase larval mortality, particularly at higher (increasing) coverage levels. For this study, we simulate LSM using $k=20$.

To further justify our choice of this larvicide potency, we examine what proportional reduction in the larvicide potency from k=20 will still result in elimination of adult mosquitos, and at what coverage level. Figure S5 shows the impact of larvicide potency and coverage on adult mosquito population. The bold black contour (labeled M=1.442) on both figures represents the threshold of the adult mosquito population (obtained as 0.1% of the minimal seasonal adult mosquito population in the absence of LSM) below which the population is considered eliminated. Any values of k and coverage to the right of these lines lead to elimination of the adult mosquito population. For instance, our analysis showed that when k=15.5 (a 22.5% reduction of larvicide potency from k=20) LSM achieves elimination of adult mosquitoes at 97.98% coverage when larvicide is applied in the stable patch alone (Figure S5A). However, when larvicide is applied both on the ephemeral and stable patches together through γ_(E ) and γ_(S ), a 37.5% less potent larvicide (larvicide efficacy reduction from k=20 to k=12.5) leads to elimination of adult mosquitoes at 98.99% coverage (Figure S5B). This means that the probability of vector elimination is not only a function of coverage but also of the duration of larvicide efficacy. In our case, using a larvicide lasting less than 6 months may not always be as effective at eliminating *An. stephensi*.

Fig. S1**. Study area and habitat characteristics.** Map of Kebridehar with the location of the 90 water-holding containers that were followed from the end of the rainy season (November 2020) and until the peak of the dry season (February 2021).

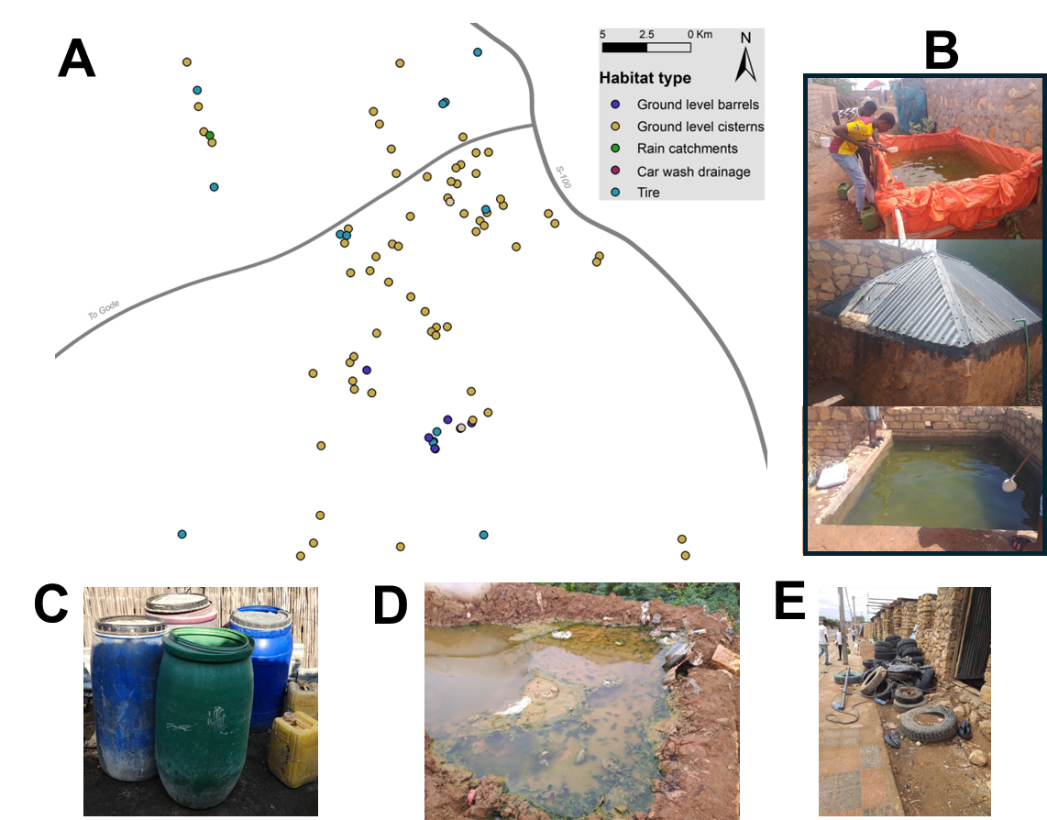

.

Fig. S2. **Temperature and rainfall for the period July 2020 to February 2021**, covering the time prior and during *An. stephensi* sampling in Kebri Dehar. Data obtained from Kebri Dehar airport weather station (code: abk).

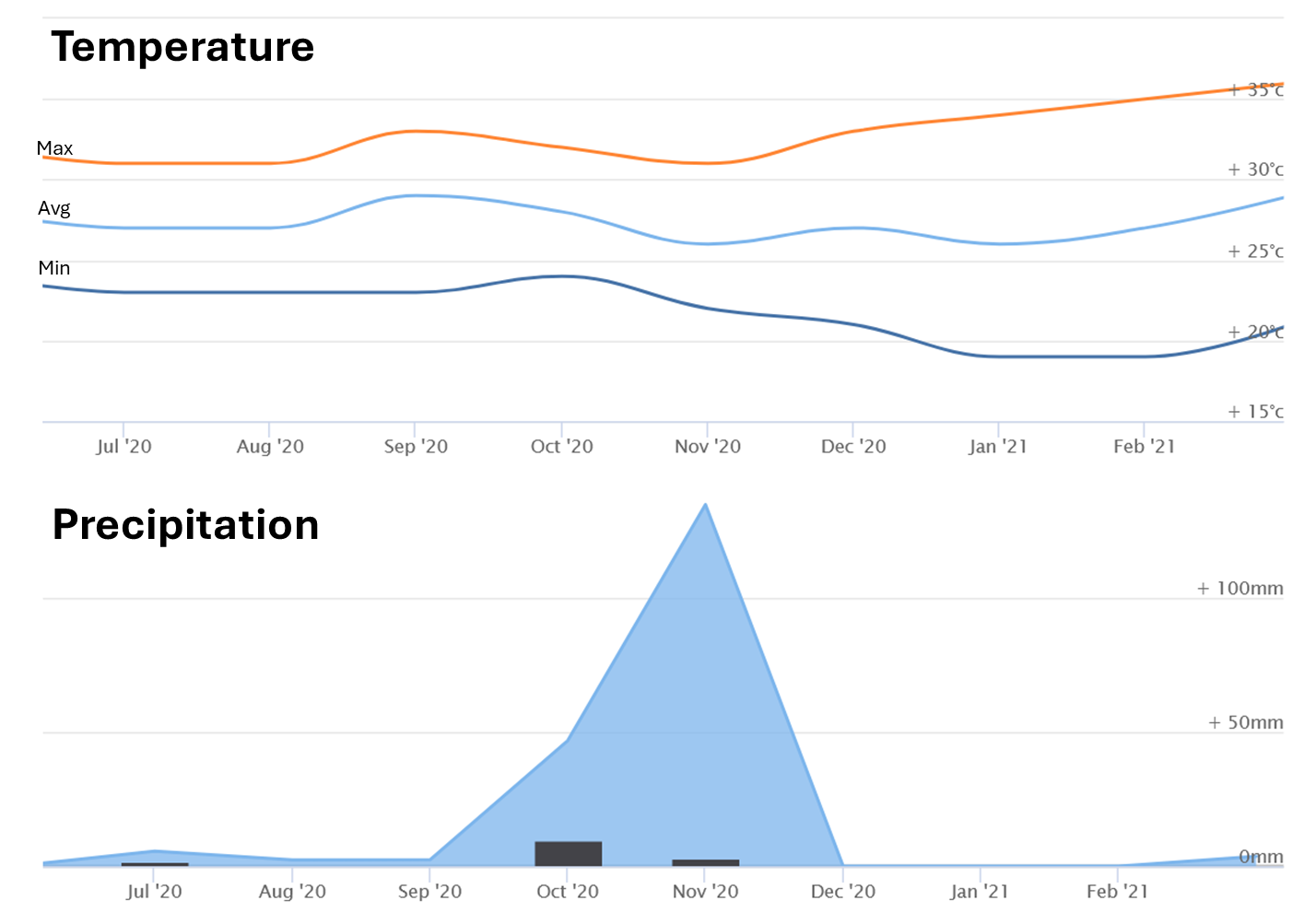

**Fig. S3. Kebri Dehar historical rainfall data.** The points are daily averages of 2010 – 2020 rainfall data, while the solid line is the smoothed version of the daily averages. The rainfall data is publicly available at https://weatherandclimate.com/ethiopia/somali/k-ebri-dehar (Accessed in July 2024).

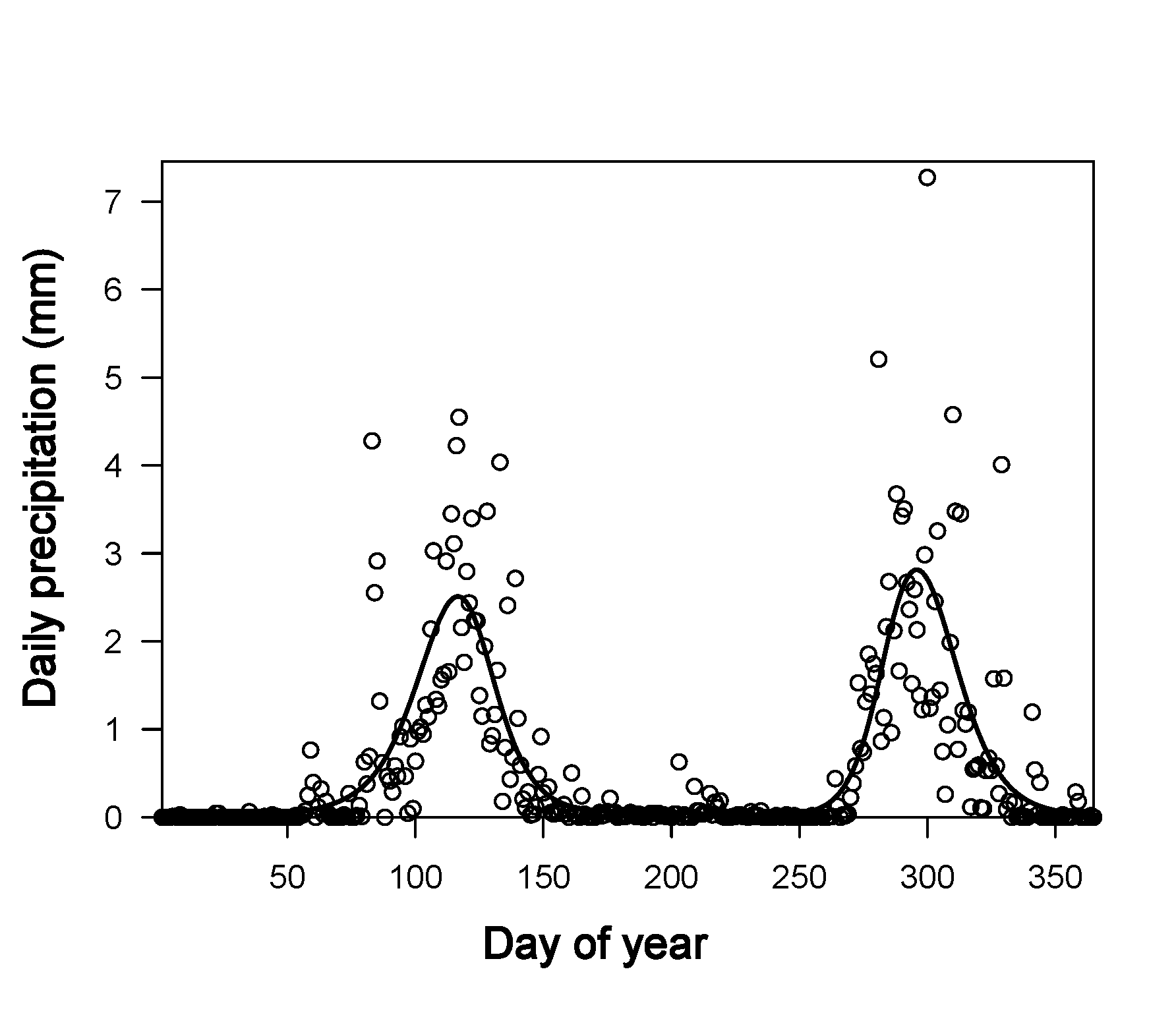

Fig. **S4. Larvae and adult *An. stephensi* mosquito population dynamics under 0% and 30% LSM coverage levels**. Impact of LSM on the entire adult population when control is performed in: a) the ephemeral patch b) the stable patch, and c) the ephemeral and stable patches together. d) Adult mosquito population (M) dynamics obtained from different coverage values. The adult mosquito population on the y-axis is the adult mosquito population value at the final simulation time for each coverage value.

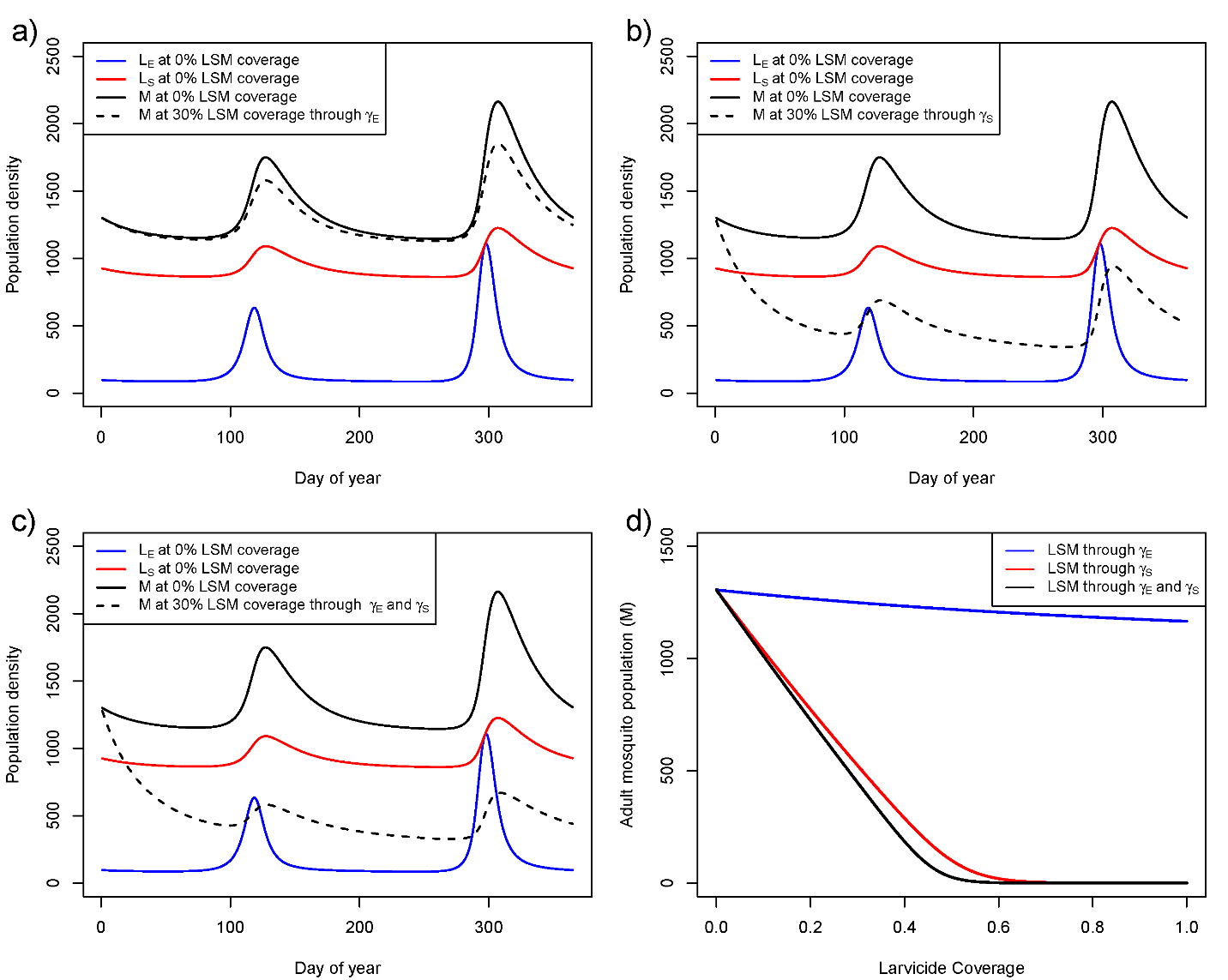

**Fig. S5. Sensitivity of LSM to the residual power of the larvicide (parameter k).** Impact of modifying the larvicide residual power (k) and the intervention coverage when LSM is implemented in the stable patch (a) and the stable and ephemeral patch (b). We consider the adult mosquito population is eliminated if their population is less than M=1.1442, a value obtained as 0.1% of the minimum of adult mosquito population in the absence of LSM.

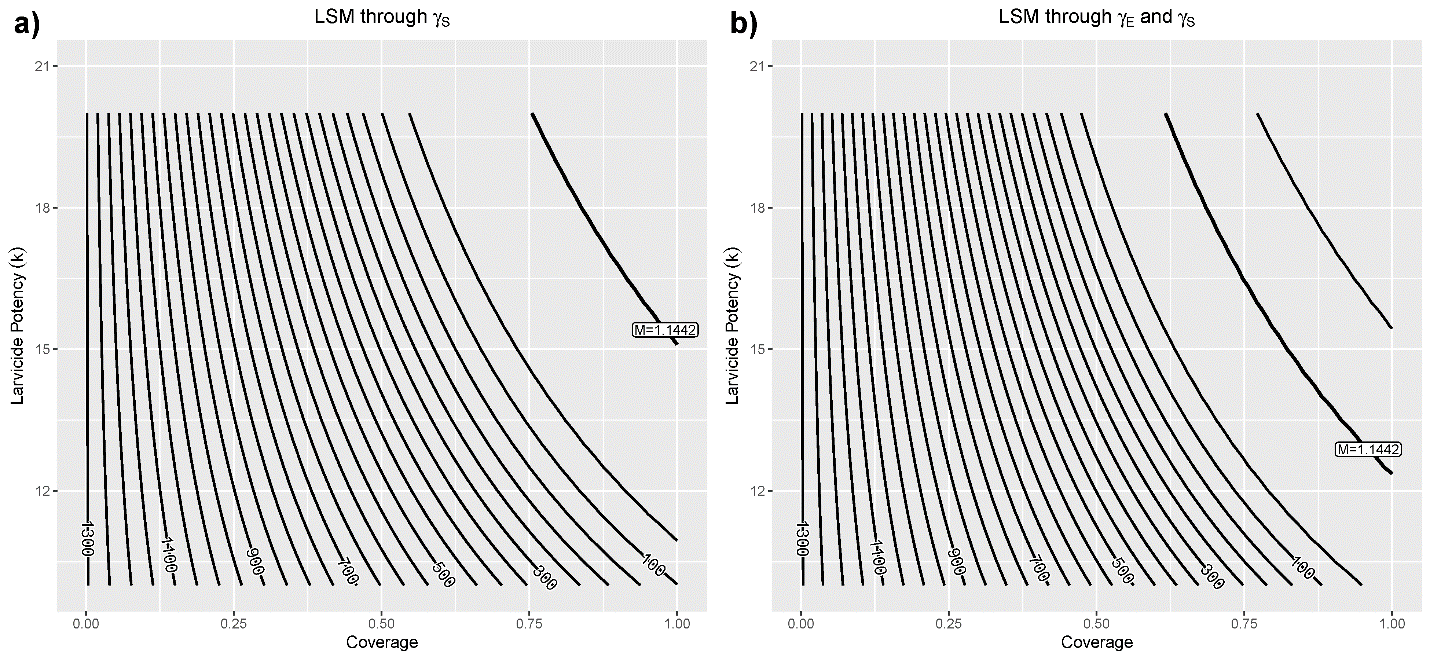

**Fig. S6.** Negative binomial fits to the distribution of *An. stephensi* larvae per container by month. Inset shows the values of the parameter *k* of the negative binomial, together with its standard deviation (SD).

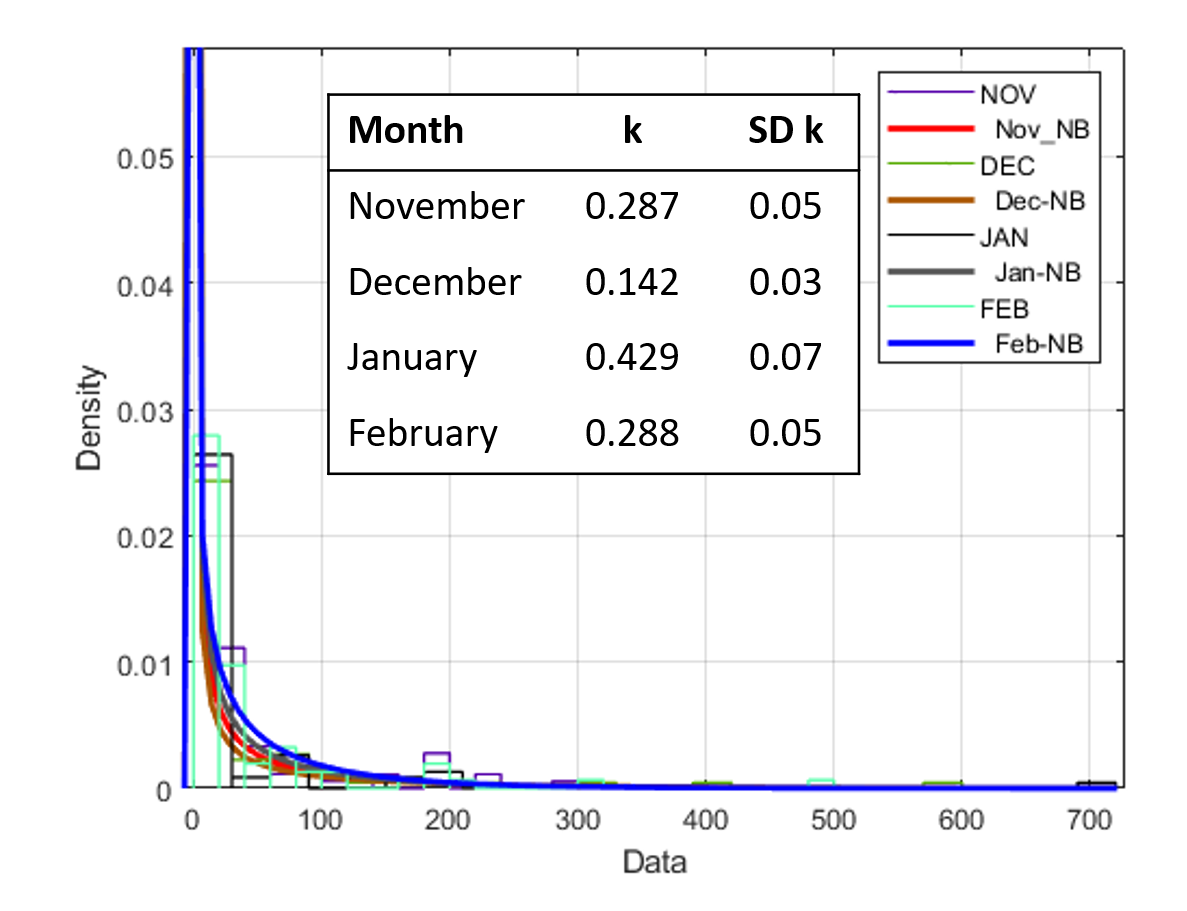

**Fig. S7.** Distribution of ‘superproductive’ habitats of *An. stephensi* in in Kebri Dehar from November 2020 to February 2021.

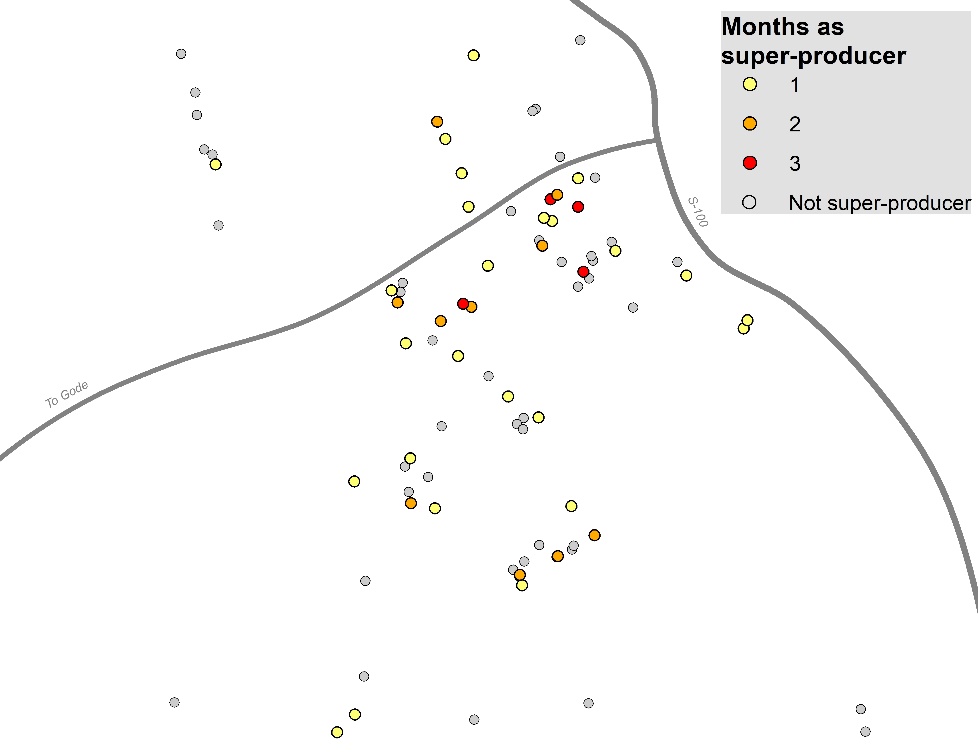
